# Deconstructing glucose-mediated catabolite repression of the *lac* operon of *Escherichia coli*: I. Inducer exclusion, by itself, cannot account for the repression

**DOI:** 10.1101/739458

**Authors:** Ritesh K. Aggarwal, Atul Narang

## Abstract

The *lac* operon of *Escherichia coli* is repressed several 100-fold in the presence of glucose. This repression has been attributed to CRP-mediated transcriptional inhibition and EIIA^Glc^-mediated inducer exclusion. The growing evidence against the first mechanism has led to the postulate that the repression is driven by inducer exclusion. The literature shows that in fully induced cells, inducer exclusion reduces the permease activity only 2-fold. However, it is conceivable that inducer exclusion drastically reduces the permease activity in partially induced cells. We measured the decline of lactose permease activity due to inducer exclusion in partially induced cells, but found that the permease activity decreased no more than 6-fold. We show that the repression is small because these experiments are performed in the presence of chloramphenicol. Indeed, when glucose is added to a culture growing on glycerol and TMG, but no chloramphenicol, *lac* is repressed 900-fold. This repression is primarily due to reversal of the positive feedback loop, i.e., the decline of the intracellular TMG level leads to a lower permease level, which reduces the intracellular TMG level even further. The repression in the absence of chloramphenicol is therefore primarily due to positive feedback, which does not exist during measurements of inducer exclusion.

## 1. Introduction

When glucose is added to a culture of *Escherichia coli* cells growing in the presence of lactose (1) or low concentrations of the gratuitous inducer TMG (2, 3), transcription of the *lac* operon is repressed several 100-fold. This so-called *catabolite repression* is usually attributed to two molecular mechanisms (4, 5):

- Inhibition of transcription mediated by 3’,5’-cyclic adenosine monophosphate (cAMP) (6, 7): Upon addition of glucose, the concentration of intracellular cAMP, and hence its complex with the cAMP receptor protein (CRP), decreases, which inhibits recruitment of RNAP to the *lac* promoter, thus reducing the *lac* transcription rate.
- Inhibition of lactose transport mediated by EIIA^glc^ (8–10): The uptake of glucose is accompanied by the formation of the dephosphorylated enzyme EIIA^glc^, which inactivates lactose permease (LacY) by binding to it, thus inhibiting lactose transport. This mechanism is called *inducer exclusion*.

Both mechanisms are well understood at the molecular level, but their contribution to catabolite repression remains the subject of debate.

Some early studies indicated that the cAMP-mediated mechanism, by itself, cannot account for catabolite repression (11, 12). However, the most compelling evidence was obtained subsequently by Aiba and co-workers who showed that the intracellular cAMP levels during the first and second exponential growth phases of the glucose-lactose diauxie were nearly the same, addition of excess cAMP did not relieve the catabolite repression of a culture growing on glucose and lactose, and catabolite repression persisted in mutants with cAMP-independent *lac* expression (1).

The foregoing evidence against the role of cAMP in catabolite repression led some researchers to posit that catabolite repression is primarily due to inducer exclusion (1, 4). This hypothesis, referred to henceforth as the *inducer exclusion hypothesis*, was supported by the following experiments. Catabolite repression does not occur if glucose is added to (a) wild-type cells growing on lactose (1) or succinate (2, 3) and *high* concentrations of a gratuitous inducer, and (b) *lacI*^-^ cells growing on lactose (1) or succinate (13). In these cases, one can argue that inducer exclusion occurred (i.e., LacY was inactivated), but was ineffective in mediating *lac* repression because the repressor was inactivated by LacY-independent mechanisms. Indeed, entry of gratuitous inducers at high concentrations is mediated primarily by passive diffusion rather than LacY (14, 15), and *lacI*^*-*^ cells synthesize defective Lac repressor that cannot bind to the *lac* operator. Catabolite repression also does not occur if glucose is added to (c) *crr*^*-*^ cells, which lack functional EIIA^glc^, or (d) *lacY*-overexpressing cells, growing on lactose (16). In these cases, it can be argued that inducer exclusion does not even exist because the EIIA^glc^:LacY ratio is so small that most of the LacY molecules escape EIIA^glc^ binding. Taken together, these data suggest that inducer exclusion is *necessary* for catabolite repression since rendering it ineffective or non-existent abolishes catabolite repression.

Although inducer exclusion may be necessary for catabolite repression, it is *not sufficient* because there is hardly any repression even if inducer exclusion exists and is effective, e.g., in the presence of active repressor and large EIIA^glc^:LacY ratios. This conclusion follows from direct measurements of the magnitude of inducer exclusion *φ*_*p*_, which is defined as the fractional reduction of LacY activity in the presence of glucose

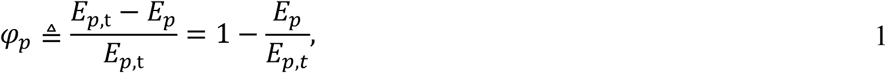

where *E*_*p*_ and *E*_*p*,t_ are the specific permease activities in the presence and absence of glucose, respectively. The determination of *φ*_*p*_ is based on the following fact: If the cells are exposed to a sufficiently small concentration of the gratuitous inducer TMG and the protein synthesis inhibitor chloramphenicol, the specific permease activity *E*_*p*_ remains constant, and the concentration of intracellular TMG, denoted *T*, approaches a steady state value 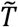 that is proportional to the specific permease activity *E*_*p*_ (Supplement Section S1). Under these conditions, Eq. 1 implies that *φ*_*p*_ is given by the expression

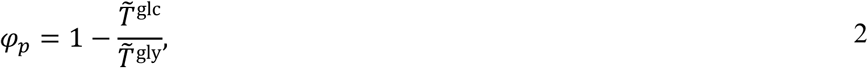

where 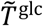 and 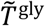 denote the steady state concentrations of intracellular TMG in the presence of glucose and a non-PTS substrate such as glycerol. Henceforth, we shall refer to these experiments for determining *φ*_*p*_ as *inducer exclusion experiments*. With the help of such experiments, Mitchell et al. determined *φ*_*p*_ at various EIIA^glc^:LacY ratios (17). They found that when the EIIA^glc^:LacY ratio was small, there was no inducer exclusion (*φ*_*p*_ *=* 0), which is consistent with the foregoing explanation of the data for *crr*^*-*^ and *lacY*-overexpressing cells. As expected, *φ*_*p*_ increased with the EIIA^glc^:LacY ratio, but it plateaued at a maximum value of only 0.5. Thus, only half of the LacY molecules were inactivated by inducer exclusion even when the repressor was active, and the concentration of EIIA^glc^ was comparable to, or higher than, the concentration of LacY. It follows that inducer exclusion, by itself, cannot account for the 600-fold catabolite repression.

The foregoing data of Mitchell et al. show that inducer exclusion is not sufficient for catabolite repression, but upon closer inspection they reveal two issues that could give us pause. First, the extracellular TMG concentrations used in their inducer exclusion experiments were so large that the steady state intracellular TMG concentrations were almost independent of the permease level (rather than proportional to it). Under this condition, *φ*_*p*_ is underestimated because similar intracellular TMG levels are obtained in the presence of glucose and glycerol 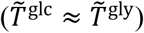 despite rather different permease levels (Supplement Section S1). Second, they varied the EIIA^glc^:LacY ratio in a non-physiological manner. Indeed, the intracellular EIIA^glc^ levels are constant (18), and the LacY level changes depending on the *lac* induction level. However, in their experiments the level of LacY, expressed from a *lac*-constitutive promoter, was constant and the EIIA^glc^ level was varied.

We were therefore led to revisit the experiments of Mitchell et al, i.e., we determined the variation of *φ*_*p*_ with the EIIA^glc^:LacY ratio, but under conditions that eliminated the two issues mentioned above. To this end, we exposed wild-type *E. coli* to a relatively low extracellular TMG level (2.5 μM) which ensured that the steady state intracellular TMG concentration was proportional to the permease level, and we varied the EIIA^glc^:LacY ratio by changing the levels of LacY rather than EIIA^glc^. Although our *φ*_*p*_ values are significantly higher than those of Mitchell et al, they are still too small to account for catabolite repression. Indeed, in fully induced cells, only half of the LacY molecules were inactivated in the presence of glucose. This fraction increased in partially induced cells (characterized by higher EIIA^glc^: LacY ratios), but it plateaued at a maximum value of 0.84, which corresponds to 1⁄0.16 ≈ 6-fold repression.

It is therefore clear that is a vast discrepancy between the repression due to inducer exclusion and catabolite repression. Indeed, if glucose is added to cells growing in the presence of low concentrations of TMG *and chloramphenicol*, the LacY activity is reduced no more than 6-fold. In contrast, if glucose is added to cells growing in the presence of low concentrations of TMG, *but no chloramphenicol*, the LacZ activity is reduced several 100-fold. It follows that inducer exclusion, by itself, cannot account for glucose-mediated catabolite repression — there is some other process that amplifies the 6-fold repression several 100-fold, which is missing from the inducer exclusion experiments.

It is useful to identify this missing process, which can be done by observing the evolution of the intracellular TMG and LacZ levels upon addition of glucose to a culture growing in the presence of low concentrations of TMG, but no chloramphenicol. We find that if glucose is added to a culture growing on glycerol and 100 µM TMG but no chloramphenicol, *lac* expression ultimately decreases 900-fold. The bulk of this repression is driven by the autocatalytic enzyme dynamics unleashed in the absence of chloramphenicol. Indeed, the decline of the intracellular TMG level leads to a decline of the permease level, which results in even lower intracellular TMG levels, and the relentless repetition of this cycle drives the permease and intracellular TMG to vanishingly low levels. Thus, enzyme synthesis, which generates positive feedback, is the process missing from the inducer exclusion experiments. The abolition of this process in the presence of chloramphenicol leads to the low repression observed in the inducer exclusion experiments. We show that the abolition of positive feedback is also responsible for the low repression observed upon addition of glucose to *lacI*^*-*^ cells, or wild-type cells exposed to *high* concentrations of TMG. In the sequel to this paper (19), we show that positive feedback plays a similar role in the massive repression observed upon addition of glucose to a culture growing in the presence of lactose rather than low concentrations of TMG.

## 2. Materials and Methods

### 2.1 Chemicals

Reagent-grade glycerol, glucose, isopropyl β-D-thiogalactopyranoside (IPTG) and *ortho*-nitrophenyl-*β*-galactoside (ONPG) were obtained from Fisher Scientific; methyl-*β*-D-thiogalactoside (TMG) was obtained from Sigma-Aldrich; and ^14^C-methyl-*β*-D-thiogalactoside ([^14^C]TMG) was procured from American Radiolabeled Chemicals, Inc. (ARC 0322).

### 2.2 Bacterial strains

Experimental strains were procured from CGSC, Yale University. Wild-type *E. coli* K12 strain MG1655 (CGSC# 6300) was used for studies pertaining to the wild-type *lac* operon. We isolated Lac permease deficient (cryptic) strain RA-Y66 by UV mutagenesis and ampicillin selection of the wild-type strain MG1655 (20). The procedure is outlined below.

MG1655 cells were irradiated with UV, killing more than 99.9 % of cells from mid-exponential growth phase, followed by overnight growth in Luria-Bertani (LB) broth in dark. Cells were subcultured in the morning in M9 minimal medium supplemented with 0.4 % lactose. Ampicillin (50 μg ml^-1^) was added to the medium and cells were allowed to grow for about two generations. This enriched the cells incapable of utilizing lactose. Cells were spun and resuspended in LB broth for a second round of ampicillin selection next day. Dilutions of enriched cultures were plated on minimal agar plates containing 0.4% glycerol, 0.1 mM IPTG, and 0.003 % 5-bromo-4-chloro-3-indolyl-β-D-galactopyranoside (X-Gal). White or pale blue isolates were grown on glycerol in the absence and presence of 0.5 mM IPTG to measure the Lac permease and β-galactosidase activity by Miller assays (20). Strain RA-Y66 was identified as one of the cultures resulting in wild-type LacZ activity and lacking LacY activity in both culturing conditions. This strain was inducible with IPTG and exhibited ONPG and TMG uptake kinetics similar to the available *lac*^-^ strain JW4081 (CGSC# 10931).

### 2.3 Culturing conditions

Cells were grown at 37 °C (unless stated otherwise) in M9 minimal medium (20) containing 0.4 % glycerol as the carbon source to ensure non-inducing and non-repressing conditions. Fully induced cells were obtained by growing the cells in the presence of 0.5 mM inducer isopropyl β-D-thiogalactopyranoside (IPTG) for at least 10 generations. The cell density (gdw L^-1^) was followed by measurement of optical density at 600 nm (OD_600_) and the two were found to be related by the expression gdw L^-1^ = 0.35×OD_600_. At several time points during the mid-exponential growth phase (OD_600_>0.3), culture aliquots were sub-cultured in medium containing 0.4 % glucose and allowed to grow for 30 min to 3 hours to ensure that the glucose transport enzymes, being fully induced, exerted the maximum inducer exclusion effect. Cells were harvested at various enzyme levels by adding 50 µg ml^-1^ chloramphenicol, after which the cells were spun and resuspended in M9 minimal medium containing 50 µg ml^-1^ chloramphenicol and carbon source, namely 0.4 % glycerol (absence of glucose) or 0.4 % glucose (presence of glucose) to obtain the *experimental suspensions* that were subsequently used to estimate the lactose permease and β-galactosidase levels. Chloramphenicol, which inhibits not only protein synthesis but also degradation of permease molecules (21), was added to ensure that the enzyme levels did not change during the course of the experiment. No chloramphenicol was added in experiments involving growth on glucose.

### 2.4 Assay for induction level

The *lac* induction levels of the *experimental suspensions* were determined by the β-galactosidase assay (20). The specific β-galactosidase activity was measured in Miller units. One Miller unit corresponds to about 200 nmol ONPG hydrolysed per minute per mg cell dry weight.

### 2.5 Assay for permease activity

The specific activity of lactose permease was assayed by measuring the steady state concentration of intracellular [^14^C]TMG using Miller’s method (20), but the concentration of extracellular TMG was much lower (2.5 μM). At these low extracellular TMG concentrations, the specific permease activity is directly proportional to the steady state concentration of intracellular TMG (Supplement), which in turn was measured as follows. At t=0, [^14^C]TMG with the indicated specific radioactivity and 2.0–2.5 µM final concentration was added to the *experimental suspensions* of both wild-type and cryptic strains which had been pre-equilibrated at 30 °C on a shaker-incubator. The intracellular TMG levels were found to reach steady state within 5 min of incubation (Fig. S3, supplement). Culture aliquots of 1 ml were therefore rapidly filtered and washed with 3 ml of medium after about 10 minutes of incubation at 30 °C. The filters were dried in scintillation vials, resuspended in a scintillation cocktail (22), and the counts were measured with a liquid scintillation counter (Perkin-Elmer Tri-Carb 2810) for at least 30 minutes to minimize the counting errors. Cellular suspensions containing no radioactivity, and M9 medium containing only [^14^C]TMG were also filtered and counted. The observed cpm were obtained after subtracting the background counts and the counts due to non-specific binding of TMG to the filter. The intracellular concentrations were calculated by assuming that the cellular water volume was 2.7 ml gdw^-1^ (23).

## 3. Results

### 3.1 Eliminating the effect of permease-mediated efflux and non-specific influx

Since our method for determining *φ*_*p*_ rests upon the validity of the linear relation 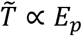, it is essential to eliminate all sources of disproportionality.

The first source of disproportionality is permease-mediated efflux of TMG which can become significant in highly induced cells due to the accumulation of large intracellular TMG levels. This phenomenon has the effect of making 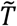 independent of *E*_*p*_, but it can be eliminated by performing the experiments with sufficiently small concentrations of extracellular TMG, thus preventing the accumulation of large intracellular TMG concentrations (Supplement Section S1). In all our experiments, the extracellular TMG concentration was 2.5 μM. Under this condition, the intracellular TMG concentration never exceeded 0.3 mM, which is negligible compared to the reported saturation constant of 84 mM for permease-mediated efflux (24). It is therefore expected that permease-mediated efflux will be negligible.

The second source of disproportionality is non-specific inducer transport mediated by diffusion or permeases other than lactose permease, which is significant in non-induced and partially induced cells (Supplement Section S1). To ensure that 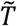 remains proportional to *E*_*p*_, it is necessary to determine the permease-mediated inducer accumulation by subtracting the non-specific inducer accumulation from the total inducer accumulation. We quantified the non-specific inducer accumulation by measuring the steady-state intracellular TMG concentration attained in the cryptic (*lacY*^*-*^) strain RA-Y66 (Fig. 1). In non-induced cryptic cells, the intracellular TMG concentration was essentially identical to the extracellular TMG concentration of 2.5 µM, which is expected since in cryptic cells, TMG is transported entirely by diffusion. In induced cryptic cells, the intracellular TMG concentration was slightly higher than the extracellular TMG concentration possibly because strain RA-Y66 is not perfectly cryptic. Importantly, the intracellular TMG concentration in non-induced wild-type cells was also comparable to the extracellular TMG concentration, which implies that in such cells also, TMG is transported almost entirely by diffusion. Hence, the determination of permease-mediated TMG accumulation is prone to significant error because it is calculated as the difference between total and non-specific TMG accumulation, but both quantities have very similar magnitudes under these conditions. In fact, we found that the total TMG accumulation was well above the non-specific TMG accumulation only in cells induced to >2 % of fully induced levels, and hence, our data are precise only for such cells.

**Fig. 1.**
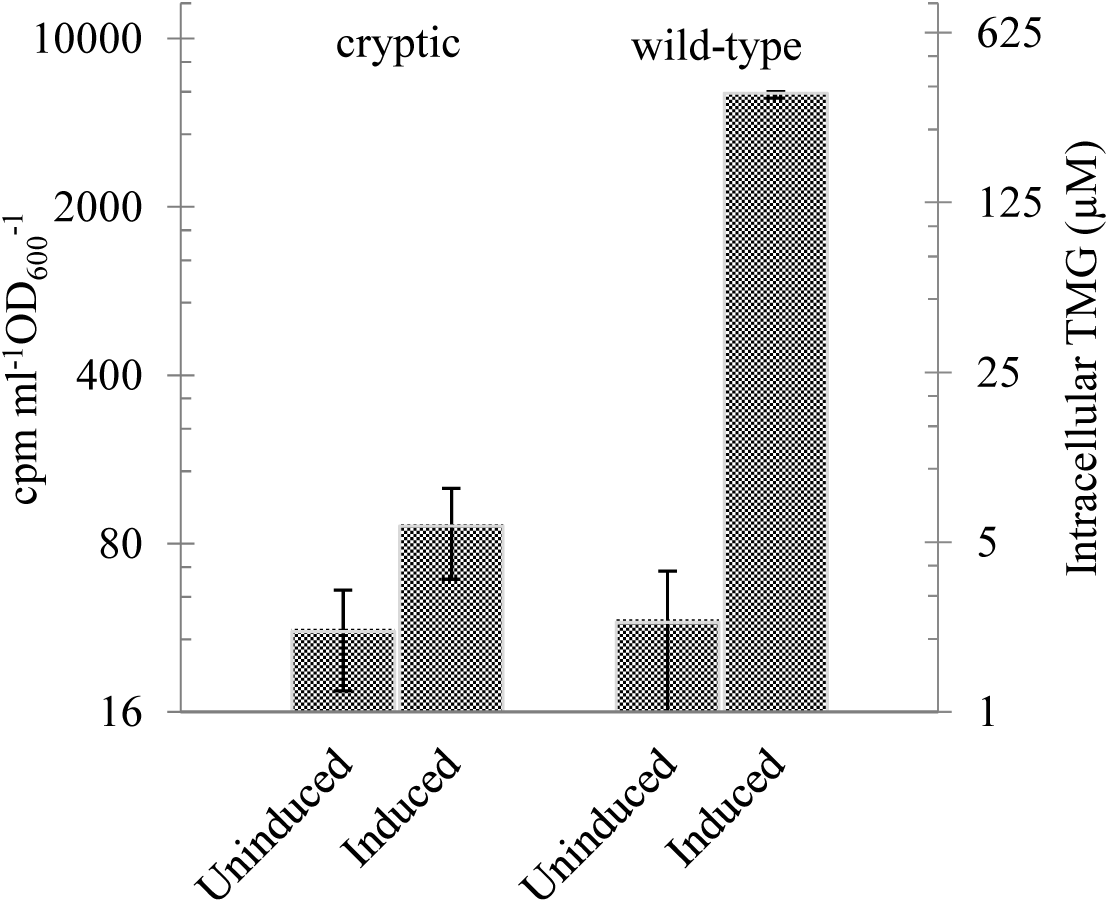
Comparison of steady state intracellular [^14^C]TMG levels in non-induced and induced cryptic (*lacY*^-^) and wild-type cells. The cpm ml^-1^ OD_600_^-1^ reported were corrected for background activity and [^14^C]TMG non-specifically bound to the filter. The intracellular concentrations on the secondary y-axis were calculated from the cpm ml^-1^ OD_600_^-1^ as described in Materials & Methods.

We confirmed that in cells induced from 2–100 % of fully induced levels, the steady state intracellular TMG concentration 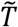, suitably corrected for non-specific accumulation, was proportional to the specific permease activity *E*_*p*_. Indeed, 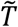 is proportional to *E*_*p*_, i.e., 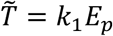 precisely if the steady state intracellular TMG concentration in the presence of glycerol is proportional to the specific β-galactosidase activity *E*_*g*_. To see this, observe that since the permease and β-galactosidase are synthesized co-ordinately, the total concentration of permease is proportional to the concentration of β-galactosidase, i.e, *E*_*p,t*_ *= k*_2_*E*_*g*_. In the presence of inducer exclusion, a fraction of the permease molecules are inactivated so that

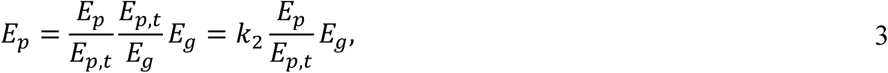

and it follows that

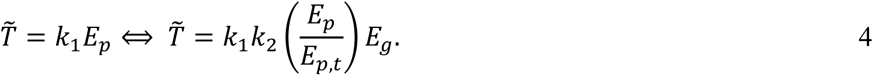

Now in the presence of glycerol, there is no inducer exclusion (*E*_*p*_ *= E*_*p,t*_), and Eq. 4 says that 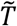 is proportional to *E*_*p*_ precisely if 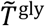 is proportional to *E*_*g*_. Fig. 2A confirms that 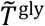 is indeed proportional to *E*_*g*_. We are therefore assured that the sources of disproportionality were successfully eliminated, and the steady state TMG accumulation 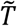 is proportional to the specific permease activity *E*_*p*_.

**Fig. 2.**
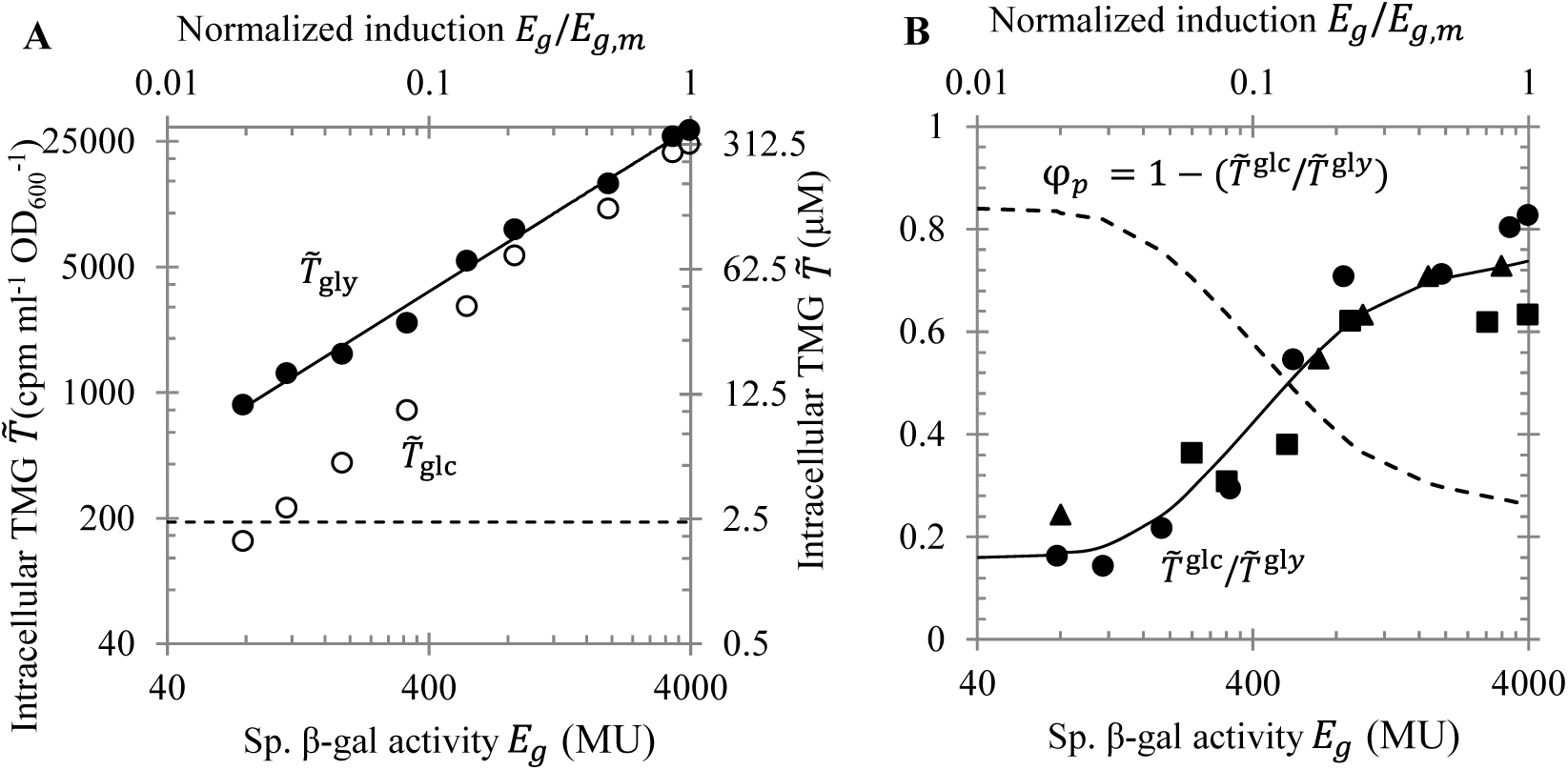
Variation of the steady state intracellular [14C]TMG concentration and magnitude of inducer exclusion with the *lac* induction level, as measured by the specific β-galactosidase activity *E*_*g*_ or the normalized induction level *E*_*g*_/*E*_*g,m*_ where *E*_*g,m*_ is the specific β-galactosidase activity of fully induced cells. (A) When the *lac* induction level is increased, the steady state intracellular [14C]TMG concentration attained in the presence of glycerol 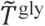 increases linearly (●), while that attained in the presence of glucose 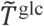 increases nonlinearly (○). The intracellular [14C]TMG concentrations were obtained by subtracting the average measured in cryptic cells. (B) When the *lac* induction level is increased, the ratio of steady state intracellular [14C]TMG levels 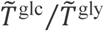 increases, but it plateaus at both large and small induction levels. The full curve shows the fit to the data obtained in three replicate experiments (solid symbols). The dashed curve shows the magnitude of inducer exclusion 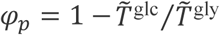 calculated from the fit to the data for 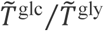.

### 3.2 The magnitude of inducer exclusion has a weak dependence on the induction level

Since 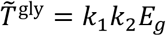, and *E*_*p*_ *< E*_*p,t*_ in the presence of glucose, Eq. 4 yields

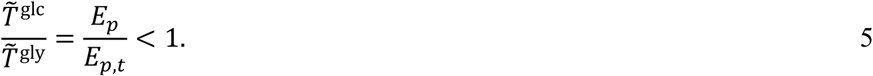

Fig. 2A shows that as expected, 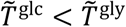 at all induction levels. The full curve of Fig. 2B shows that the ratio 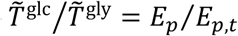 increases with the induction level *E*_*g*_. In fully induced cells, this ratio is 0.80, but still increasing, and likely to become 1.0 in hyper-induced cells. Moreover, the lower the induction level, the lower the ratio 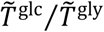, but it plateaus at 0.16, which corresponds to a mere 100/16 ≈ 6-fold reduction of the specific permease activity. Equivalently, the dashed curve Fig. 2B shows that the magnitude of inducer exclusion *φ*_*p*_ *=* 1 − *E*_*p*_⁄*E*_*p,t*_ decreases with the induction level, but it never exceeds 0.84.

The limiting values of *E*_*p*_⁄*E*_*p,t*_, equalling 1.0 in hyper-induced cells and 0.16 in hypo-induced cells, are consistent with the limiting values obtained from a simple stoichiometric binding model of inducer exclusion. To see this, observe that before glucose is added to the experimental suspension, all the EIIA^glc^ molecules are phosphorylated and all permease molecules are active, i.e., not bound to EIIA^glc^. Let 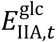 and *E*_*p,t*_ denote the intracellular concentrations of phosphorylated EIIA^glc^ and permease under these conditions. Upon addition of glucose, virtually all the EIIA^glc^ molecules are dephosphorylated (25), and some of them bind to the free permease molecules in the stoichiometric ratio 1:1 (26, 27), thus forming the inactive complex 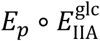, i.e., the binding can therefore be represented by the reaction

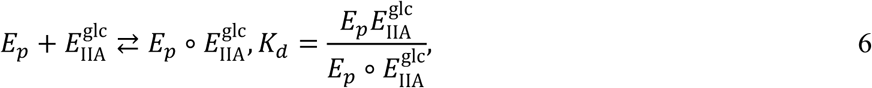

where *K*_*d*_ is the dissociation constant. Now in hyper-induced cells, the permease molecules are in excess, i.e., 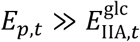. Hence the fraction of permease molecules inactivated by binding is negligibly small, i.e., *E*_*p*_ ≈ *E*_*p,t*_ and 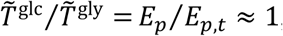, which is consistent with the data for hyper-induced cells obtained by Mitchell et al. In hypo-induced cells, 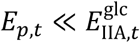 which implies that the fraction of EIIA^Glc^ molecules bound to permease is negligibly small, i.e., 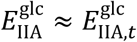. Thus, the binding reaction reduces to the pseudo-first-order reaction 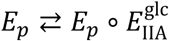 with dissociation constant 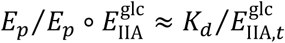, which implies that

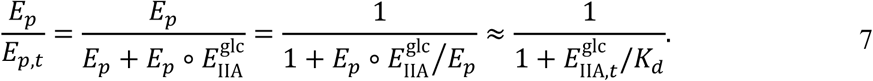

Since 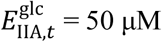 and *K*_*d*_ = 10±5 μM (27), we obtain *E*_*p*_/*E*_*p,t*_ = 1/6, which is consistent with our data for hypo-induced cells shown in Fig. 2B. Thus, even in the presence of active repressor and large EIIA^glc^:LacY ratios, inducer exclusion yields no more than 6-fold repression. It seems desirable to understand why the repression due to inducer exclusion is so small.

### 3.3 Strong catabolite repression can occur even in the presence of TMG

We show below that the repression observed in the inducer exclusion experiments is small because enzyme synthesis is abolished in the presence of chloramphenicol. If these experiments are performed in the absence of chloramphenicol, the underlying enzyme dynamics are unleashed and give rise to strong repression. More precisely, the addition of glucose to a culture growing on glycerol and TMG, but no chloramphenicol, leads to a dramatic reduction in *lac* expression.

We can infer the potential for strong glucose-mediated catabolite repression in the presence of TMG by examining the extensive data obtained by Ozbudak et al. (Fig. 3). They showed that if *E. coli* cultures were exposed to the non-repressing carbon source succinate and various concentrations of extracellular TMG, but no chloramphenicol, bistability occurred when the extracellular TMG concentration was in the range 3–30 µM. Bistability also occurred if similar experiments were performed with a mixture of succinate and the repressing carbon source glucose, but the bistable region was now shifted to a higher range of extracellular TMG concentrations (100–1000 µM). These data imply that if glucose is added to a culture growing exponentially on succinate and 30–100 µM TMG, the specific β-galactosidase activity will ultimately decrease several 100-fold.

**Fig. 3.**
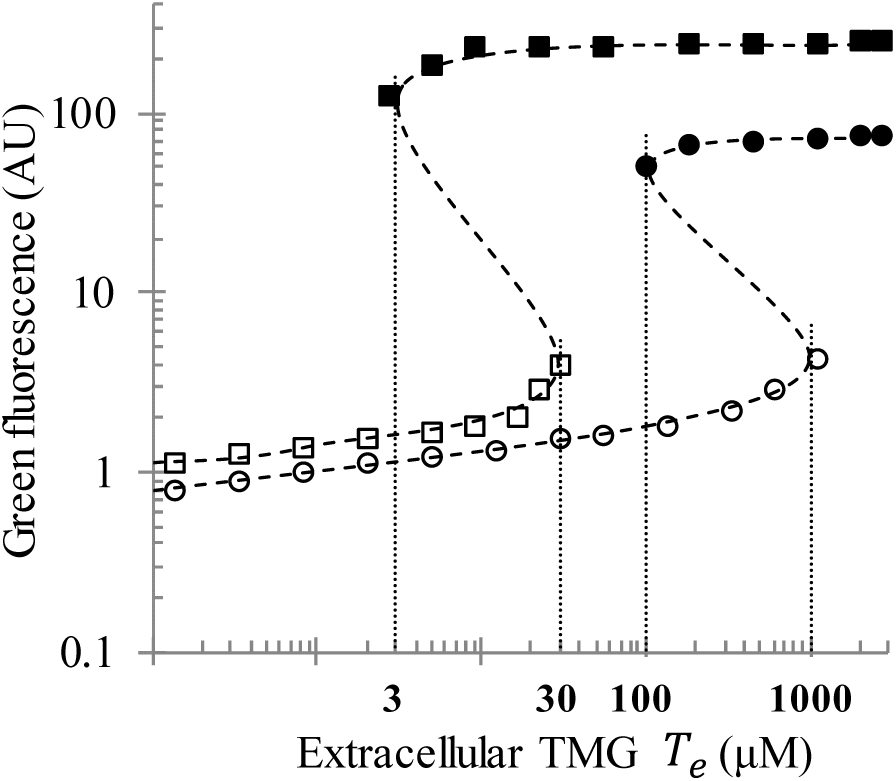
Schematic diagram showing the steady state *lac* promoter activity, as measured by the green fluorescence intensity, during growth of *E. coli* cultures on succinate (squares) and succinate + 1 mM glucose (circles) in the presence of various extracellular TMG concentrations (adapted from (3)). The open (resp., closed) symbols show the steady state GFP intensity reached when the inoculum is uninduced (resp., induced). When the extracellular TMG concentrations are 30–100 µM, cells grown on succinate are eventually induced, but cells grown on succinate + 1 mM glucose are eventually uninduced. It follows that if 1 mM glucose were added to a steady state culture growing on succinate and 30–100 µM extracellular TMG, the *lac* promoter activity should decrease several 100-fold, which is similar in magnitude to the catabolite repression observed in the glucose-lactose diauxie.

Since glycerol, like succinate, is also a non-repressing carbon source, we reasoned that data similar to Fig. 3 would be obtained if succinate were replaced by glycerol. More precisely, we hypothesized that if glucose were added to a culture growing exponentially on glycerol and certain concentrations of extracellular TMG, but no chloramphenicol, the specific β-galactosidase activity would ultimately decrease to dramatically low levels. We found that this was indeed the case when the concentration of extracellular TMG was 100 µM. When induced or non-induced cells of wild-type *E. coli* were grown overnight on glycerol and 100 µM TMG, they became fully induced (∼3000 Miller units) regardless of their initial state. In contrast, if glucose (4 g L^-1^) was added to a steady state culture growing exponentially on glycerol and 100 µM TMG, the specific β-galactosidase activity and intracellular [^14^C]TMG concentrations declined continuously from the high levels observed before the addition of glucose (Fig. 4A). Importantly, this decline was biphasic — within 5 min, the intracellular TMG level declined ∼3-fold while the specific β-galactosidase activity remained constant, but after this rapid initial transient, the intracellular TMG concentration and specific β-galactosidase activity decreased in tandem, slowly but relentlessly. Although the intracellular TMG level could not be measured precisely at low induction levels, the specific β-galactosidase activity was ultimately *<* 3.5 Miller units, which corresponds to ∼900-fold repression.

**Fig. 4.**
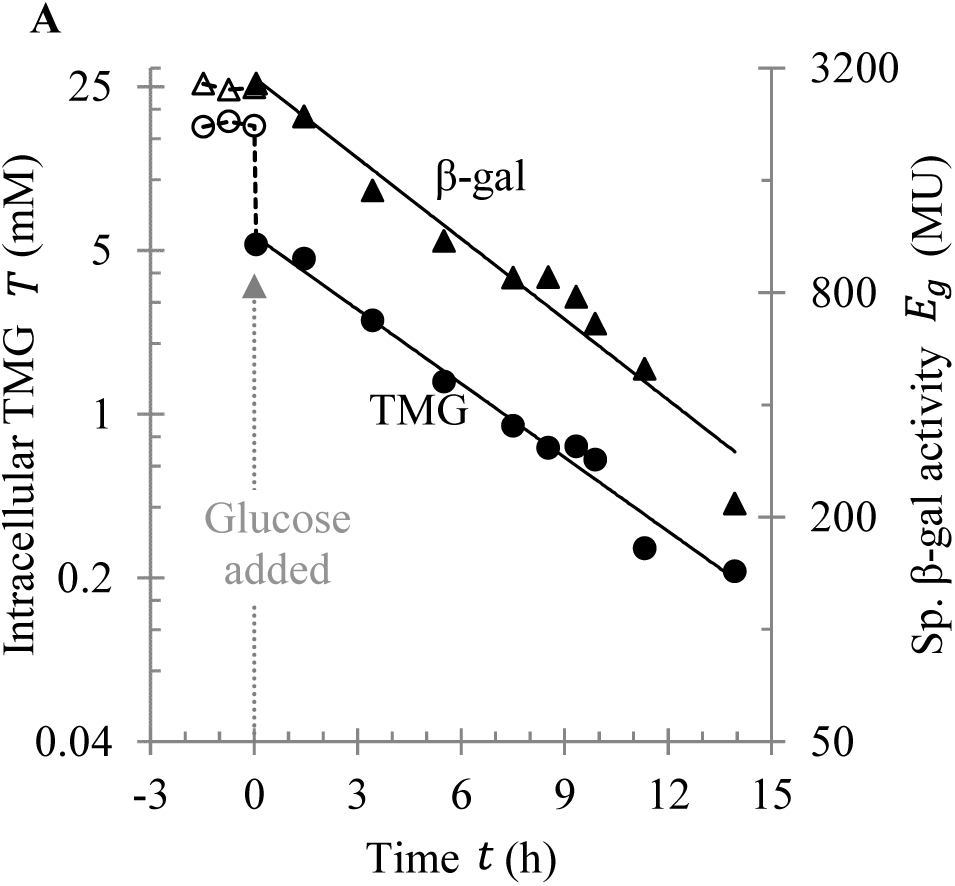
Strong catabolite repression in the presence of TMG. (A) Wild-type cells were pre-grown at 30 °C on glycerol and 100 μM [^14^C]TMG until they reached steady state, as indicated by the constant specific β-galactosidase activity (△) and intracellular TMG level (○) at *t* < 0. At *t* = 0, 0.4 % glucose was added, and the specific β-galactosidase activity (▴) and intracellular TMG level (●) were measured for 7 generations. Within 5 min of glucose addition, the intracellular TMG level decreased 3-fold, whereas the specific β-galactosidase activity remained unchanged. Thereafter, both slowly decreased approximately 15-fold over the next 15 h.

The initial decline of the intracellular TMG level, which occurs while the β-galactosidase level is essentially constant, is probably due to inducer exclusion since its magnitude (3-fold) and time scale (5 min) are similar to those observed in the inducer exclusion experiments (Fig. 2A and Fig. S3). We show below that the subsequent slow and simultaneous decline of the intracellular TMG and β-galactosidase levels, which is not observed in the inducer exclusion experiments of Fig. 2A, is primarily due to the slow evolution of the lactose enzymes unleashed in the absence of chloramphenicol.

### 3.4 Proposed kinetics of catabolite repression in the presence of TMG

Before proposing a kinetic model for the strong glucose-mediated repression observed in Fig. 4, it is useful to recall two key facts.

First, our earlier experiments showed that the steady state intracellular TMG concentrations 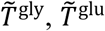 are increasing functions of the induction level *E*_*g*_, linear in the case of glycerol and non-linear in the case of glucose (Fig. 2A). Although these data were obtained at various *fixed E*_*g*_, engineered by the addition of chloramphenicol, we expect similar data if the induction level *E*_*g*_ changes *quasi-statically* due to the slow evolution of the enzymes unleashed in the absence of chloramphenicol (Supplement Section S2). Thus, based on the data in Fig. 2A, the variation of the quasi-steady state intracellular TMG levels, 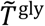 and 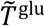, with the slowly varying induction level *E*_*g*_ can be depicted graphically by the dashed linear line and full nonlinear curve in Fig. 5A.

**Fig. 5.**
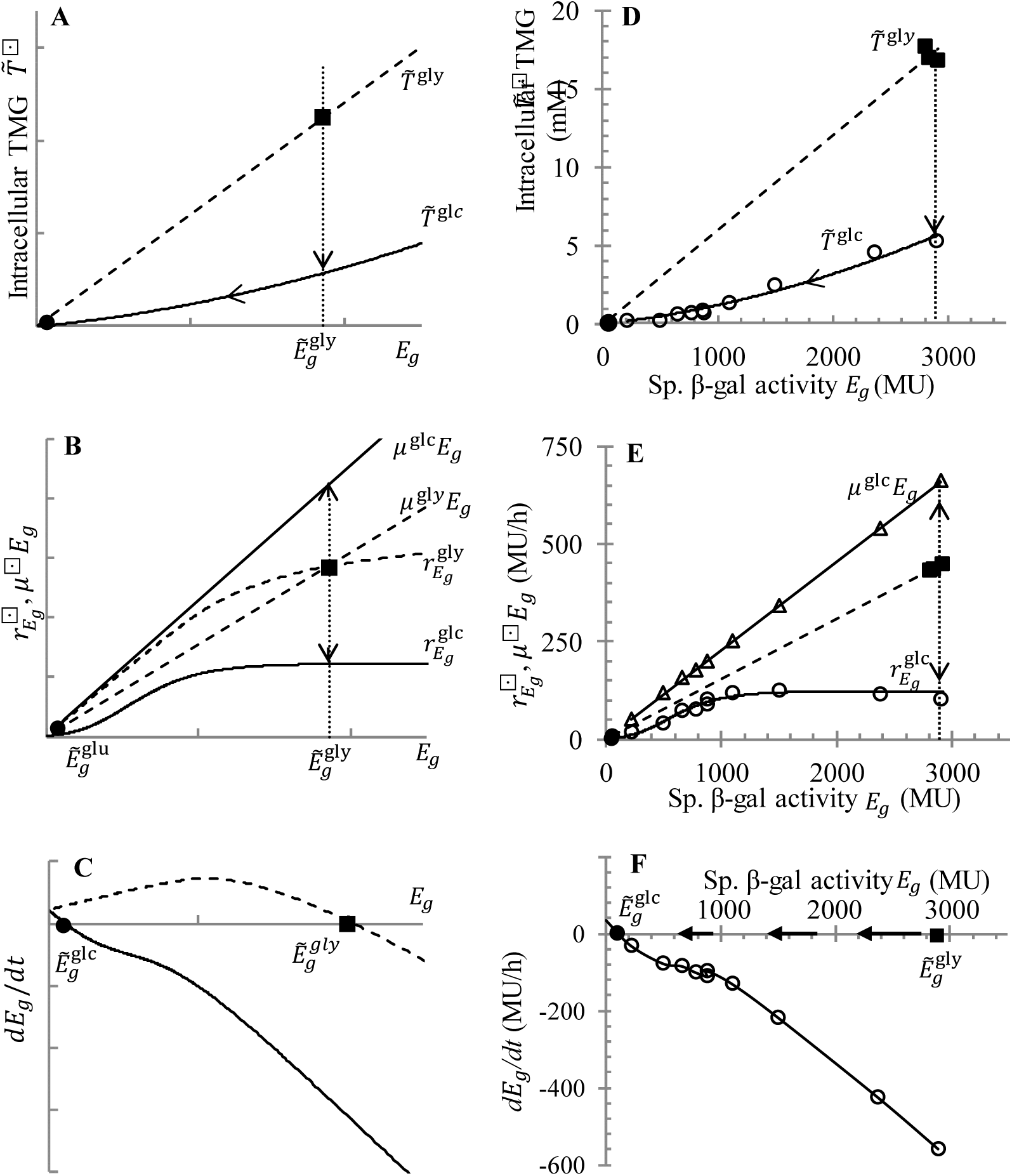
Proposed model for the glucose-mediated catabolite repression shown in Fig. 4 and its comparison with data. *Left panel:* Hypothesized variation with the induction level *E*_*g*_ of (A) the quasi-steady state intracellular TMG levels 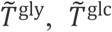, (B) the specific rates of β-galactosidase synthesis 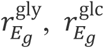 and dilution *µ*^gly^*E*_*g*_, *µ*^gly^*E*_*g*_, and (C) the net rate of β-galactosidase synthesis *dE*_*g*_/*dt*. The dashed and solid curves show these variations during growth of *E. coli* in the presence of glycerol + TMG and glucose + glycerol + TMG, respectively. *Right panel:* Observed variations with induction level *E*_*g*_ of (D) the intracellular TMG level 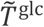, (E) the rates of induction 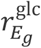 and dilution *µ*^gly^*E*_*g*_,, and (F) the rate of change of induction level *dE*_*g*_/*dt*. These variations are derived from the data in Fig. 4.

Second, since β-galactosidase is relatively stable (28), the slow evolution of the induction level enzyme *E*_*g*_ is governed by the rates of β-galactosidase synthesis and dilution due to growth, i.e.,

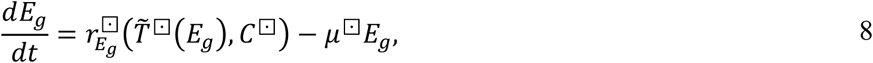

where the superscript ⊡ specifies the experimental condition (⊡*=* gly when the experiment is performed with glycerol + TMG; ⊡*=* glc when the experiment is performed with glucose + glycerol + TMG), *C*^⊡^ denotes the intracellular cAMP level at the given experimental condition, and 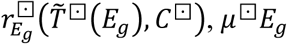 denote the induction and dilution rates of β-galactosidase at the given experimental condition. Since 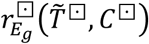 is an increasing function of the intracellular TMG level 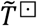 (25), and 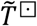 is an increasing function of *E*_*g*_ (Fig. 2A), 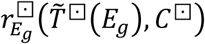 is an increasing function of *E*_*g*_, i.e., induction follows autocatalytic kinetics. If *µ*^⊡^ is constant, as was found to be the case in our experiments, the dilution rate follows first-order kinetics.

We begin by explaining *lac induction* in the presence of glycerol + 100 µM TMG. We assume that under these conditions, the synthesis rate 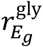 and the dilution rate *µ*^gly^*E*_*g*_ have the graphs shown in Fig. 5B as the dashed sigmoidal curve and dashed line, respectively. The disposition of the two graphs is such that they intersect at a single point, denoted ◼, which represents the steady state with specific β-galactosidase activity denoted 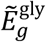; moreover, for all 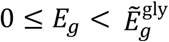, the induction rate exceeds the dilution rate, and hence, 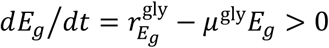 (Fig. 5C). It follows that if an uninduced culture *E*_*g*_(0) ≈ 0 is exposed to glycerol + 100 µM TMG, the induction level *E*_*g*_ increases monotonically until it reaches the steady state value 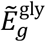.

Next, we explain the strong glucose-mediated *lac repression* observed in Fig. 4. We assume that the addition of glucose to a steady state culture growing exponentially on glycerol and 100 µM TMG has the following three effects:

1. Due to inducer exclusion, the intracellular TMG level decreases rapidly while the induction level remains essentially constant at its initial value 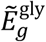, which is depicted in Fig. 5A by the downward vertical arrow from 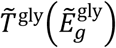 to 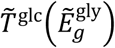. In addition, the intracellular cAMP level also rapidly declines from *C*^gly^ to *C*^glc^ (29).
2. Due to the decline of the intracellular TMG and cAMP levels, the induction rate decreases from 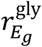 to 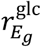, i.e., the graph of the induction rate shifts downward from the dashed to the full sigmoidal curve in Fig. 5B.
3. Due to glucose uptake and metabolism, the specific growth rate increases, i.e., the line representing the dilution rate rotates counter-clockwise from the dashed to the full line in Fig. 5B.

Due to the first effect, the intracellular TMG and cAMP levels decline, but *E*_*g*_ is essentially constant. Due to the second and third effects, the disposition of the induction and dilution curves changes such that they intersect at a new steady state, denoted ●, such that the corresponding induction level 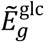 is drastically lower than the steady state induction level 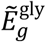 attained during growth on glycerol + TMG (Fig. 5B); moreover, for all 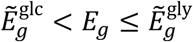, the dilution rate now exceeds the induction rate, and hence, *dE*_*g*_/*dt* < 0 (Fig. 5C). It follows that the induction level *E*_*g*_ decreases relentlessly from its initial value 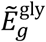 to its new steady state value 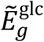. During this slow decline of *E*_*g*_, the intracellular TMG level, which is in quasi-steady state, follows the curve labelled 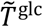in Fig. 5A.

### 3.5 Proposed kinetics of catabolite repression is consistent with observed kinetics

We can test the validity of the proposed model because we measured the evolution of the biomass, β-galactosidase, and intracellular TMG concentrations after the addition of glucose to a steady state culture growing exponentially on glycerol + TMG (Fig. 4). From these measurements, we can immediately determine the variation of the quasi-steady state intracellular TMG level 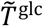 with the slowly decreasing induction level *E*_*g*_. With a little more effort, our data also yield the quasi-steady state induction and dilution *rates*, which explain why the induction level declines in the presence of glucose.

The decline of the quasi-steady state intracellular TMG level with the slowly declining induction level is consistent with the predicted trend shown in Fig. 5A. Indeed, if we plot the instantaneous intracellular TMG levels in Fig. 4 against the corresponding induction levels, we obtain the graph labelled ○ in Fig. 5D. It is clear from these data that immediately after the addition of glucose, the induction level remains unchanged at 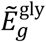, but the intracellular TMG level instantly decreases 3-fold from its previous steady state value 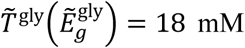, denoted the symbol ◼, to the new quasi-steady state value 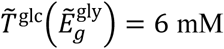, a process that is analogous to inducer exclusion (dotted vertical line with arrow). Thereafter, the quasi-steady state intracellular TMG level 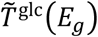 declines slowly in tandem with the induction level *E*_*g*_ (full curve with arrow joining the points labelled ○).

Next, we show that the kinetic mechanism governing the slow decline of the induction level *E*_*g*_ is also consistent with the model because the dilution and induction kinetics extracted from the data in Fig. 4 are consistent with the picture in Fig. 5B. To see this, observe that the variation of the cell density with time, measured after the addition of glucose, provides the specific growth rate *µ*^glc^, and hence the dilution rate *µ*^glc^*E*_*g*_. This dilution rate, which is derived from the data, appears in Fig. 5E as the full line passing through the origin and the points labelled Δ. Given this dilution rate *µ*^glc^*E*_*g*_, we can also determine the induction rate 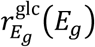 by calculating the rate of change of the specific β-galactosidase activity *dE*_*g*_/*dt* from the data, and substituting it in the relation

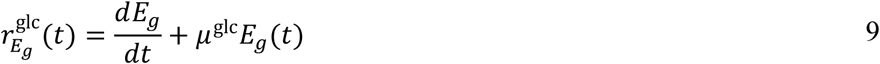

obtained from Eq. 8. When this induction rate 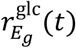, which is estimated from the data, is plotted against the corresponding induction level *E*_*g*_*(t)*, we obtain the full sigmoidal curve passing through the points labelled ○ in Fig. 5E. This sigmoidal induction rate curve intersects the dilution rate line at the point labelled ● with a very low specific β-galactosidase activity 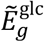 (Fig. 5E). Moreover, 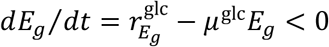 for all 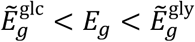 (Fig. 5F). Consequently, the induction level decreases monotonically from its initial value 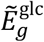, denoted ◼, to the new steady state value 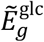, denoted ●. This leads to the persistent decline of the quasi-steady state intracellular TMG level from 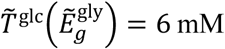 to the immeasurably small value 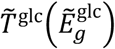.

Although the intracellular TMG level and the induction rate decrease immediately after the addition of glucose, the initial decline of the induction rate is not due to the decline of the intracellular TMG level. Indeed, Fig. 5B shows that despite the 3-fold decline of the intracellular TMG level immediately after the addition of glucose, the induction rate remains independent of the intracellular TMG level for some time, i.e., the intracellular TMG concentration is at a saturating level even after its initial decline. It follows that the initial decline of the induction rate cannot be due to inducer exclusion — it must be due to the decline of the cAMP level that occurs immediately after the addition of glucose (29). Although inducer exclusion plays no role in the initial decline of the induction rate, it must play a critical role in the subsequent decline of the induction rate when the intracellular TMG level has dropped to sub-saturating levels at which the induction rate is extremely sensitive to variations of the TMG level (30, 31).

We conclude that the enzyme dynamics unleashed in the absence of chloramphenicol bring to the fore the potent effect of positive feedback and autocatalytic induction kinetics which play a crucial role in mediating catabolite repression.

## 4. Discussion

### 4.1 Comparison of our results to studies in the literature

Our work differs from earlier work with respect to the goal as well as the method. Indeed, since the goal of earlier work to understand the mechanism of bistability, the theoretical models focused on the *steady states* (3, 32–35), and the experiments were primarily concerned with the measurement of the *induction levels* (2, 3, 36). However, our goal was to understand the mechanism of strong glucose-mediated catabolite repression, which led us to focus on the *transients* and to measure both the *induction and inducer levels*.

Our focus on the transients, and the kinetics derived from it, also provides a more conclusive confirmation of the theory. Indeed, in the earlier work cited above, the authors proposed the model kinetics, but verified the model by checking the agreement between the observed and predicted steady states. This method is not conclusive because it is conceivable that some other kinetics, distinct from those proposed in the model, also yield the same steady states. However, we acquired transient data that allowed us to calculate the induction and dilution kinetics, and these were found to be in complete agreement with the kinetics proposed in the model. In short, earlier work proposed kinetics, but verified the model by observing the steady states, whereas we proposed and verified the kinetics, which provides definitive validation of the model.

### 4.2 Relative roles of the various mechanisms that mediate catabolite repression

We have shown that upon addition of glucose to a culture growing exponentially on TMG + glycerol, the steady state induction and inducer levels decline dramatically because the induction rate decreases due to the rapid decline of the intracellular TMG and cAMP levels, and the dilution rate increases due to the enhanced specific growth rate (Fig. 5). Importantly, the magnitude of each of these effects is rather small — inducer exclusion reduces the intracellular TMG levels no more than 6-fold (Fig. 2), the decline of the cAMP levels reduces the induction rate 3-fold (Fig. 5B), and the dilution rate increases only 2-fold (Fig. 5B). Yet, these small effects, taken together, produce a drastic 900-fold repression by shifting the steady state induction level from 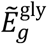to 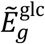.

The autocatalytic enzyme kinetics play a key role in mediating the drastic reduction of the steady state induction level — if the kinetics of enzyme synthesis are not autocatalytic, the addition of glucose produces relatively small changes of the steady state. Indeed, we have already shown that if enzyme synthesis, which is necessary for positive feedback, is abolished by adding chloramphenicol, the addition of glucose produces only a modest decline of the permease activity which reflects the effect of inducer exclusion (Fig. 2). However, the repression is massively attenuated even if enzyme is synthesized, but its synthesis rate 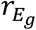 is independent of the induction level *E*_*g*_, i.e., the graph of 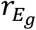 is a horizontal line rather than the increasing function shown in Fig. 5E. This can be engineered in two ways: Render 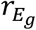 independent of 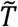, as is the case in *lac*-constitutive mutants, or make 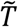 is independent of *E*_*g*_ by adding excess TMG or IPTG to the culture (in which case inducer is transported primarily by diffusion rather than by LacY). In both cases, 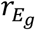 becomes independent of the induction level *E*_*g*_, and addition of glucose produces a very modest effect. Indeed, there is practically no repression if glucose is added to *lac*-constitutive mutants (13), and wild-type cells exposed to excess TMG (2). The existence of autocatalytic induction kinetics is therefore necessary for the dramatic shift of the steady state induction level.

### 4.3 Implication for the glucose-lactose diauxie of *E. coli*

We have shown that insofar as catabolite repression in the presence of the *non-metabolizable* inducer TMG is concerned, inducer exclusion, by itself, is not sufficient — the autocatalytic induction kinetics, engendered by positive feedback, play a crucial role. It is relevant to ask if this result also holds for the glucose-lactose diauxie which occurs in the presence of the *metabolizable* substrate lactose.

The insights derived from this work suggest that the answer lies in the following two questions:

1. Does positive feedback exist during growth on lactose?
2. Does positive feedback act (in the reverse direction) when glucose is added to a culture growing on lactose?

If both questions yield an affirmative answer, it is likely that the foregoing results also apply to the glucose-lactose diauxie.

Given what is known about the regulation of *lac* expression, the first question reduces to the experimentally testable question: Do the lactose enzymes (specifically, permease and β-galactosidase) promote the accumulation of allolactose? Indeed, since allolactose stimulates synthesis of the lactose enzymes, positive feedback exists precisely if the lactose enzymes promote the accumulation of allolactose, i.e., the intracellular allolactose level increases with the induction level as is the case for intracellular TMG (Fig. 5D). In the sequel to this manuscript (19), we show that this is indeed the case — the intracellular allolactose levels always increase monotonically with the β-galactosidase levels.

To address the second question, it is necessary to ask if upon addition of glucose to a culture growing on lactose, allolactose and β-galactosidase levels decline in tandem, in a manner analogous to the decline of intracellular TMG and β-galactosidase levels shown in Fig. 4. More importantly, the induction and dilution kinetics derived from the data should have the forms similar to those shown in Fig. 5E. This is also the case as shown the sequel to this manuscript (19).

The dynamics of both induction and glucose-mediated repression of the *lac* operon in the presence of lactose are therefore analogous to those observed in the presence of small concentrations of TMG.

## 5. Conclusions

We began by highlighting the discrepancy between the inducer exclusion hypothesis which postulates that the 600-fold catabolite repression is driven entirely by inducer exclusion, and the relatively small 2-fold repression reported in measurements of the magnitude of inducer exclusion. In this work, we confirmed that the magnitude of inducer exclusion is quite small with no more than 6-fold repression even in hypo-induced cells. This occurs because the experiments are performed in the presence of chloramphenicol which abolishes enzyme synthesis and destroys positive feedback. Indeed, when glucose is added to a culture growing on glycerol and small concentrations of TMG, but no chloramphenicol, the *lac* operon is repressed 900-fold and positive feedback plays a critical role in mediating this repression. The destruction of positive feedback also explains the weak repression observed upon addition of glucose to *lacI*^*-*^ cells, and wild-type cells exposed to high concentrations of TMG.

## Acknowledgement

We dedicate this work to late Prof. Frederick C. Neidhardt whose questions led to the experiments reported here. The early part of this research was supported by the DST grant SR/SO/BB-79/2010.

## ABBREVIATIONS AND NOTATIONS

**ABBREVIATIONS**

cAMP: 3’,5’-cyclic adenosine monophosphate
CGSC: Coli Genetic Stock Centre, Yale University
CRP: cAMP receptor protein
EIIA^Glc^: Enzyme III of glucose transport system, PTS (Previously EIII^Glc^)
gdw: Grams of cell dry weight, g
IPTG: Isopropyl β-D-thiogalactopyranoside or MeSH
LB: Luria-Bertani broth
MU: Miller units
OD_600_: Optical density at 600 nm
ONPG: *ortho*-nitrophenol-β-D-galactopyranoside
RNAP: Ribonucleic acid (RNA) polymerase
TMG: Methyl- β-D-1-thiogalactopyranoside
[^14^C]TMG: Carbon-14 labelled isotope of TMG
UV: Ultraviolet (radiation)
X-gal: 5-bromo-4-chloro-3-indolyl-β-D-galactopyranoside

## NOTATIONS

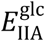: Intracellular concentration of phosphorylated EIIA^glc^ in the absence of glucose
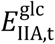: Intracellular concentration of dephosphorylated EIIA^glc^ in the presence of glucose
*E*_*g*_: Specific activity of β-galactosidase
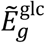: Steady state specific activity of β-galactosidase in the presence of glucose
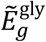: Steady state specific activity of β-galactosidase in the presence of glycerol
*E*_*p*_: Specific activity of Lac permease
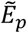: Steady state specific activity of Lac permease
*E*_*p*,t_: Specific activity of free Lac permease in the absence of glucose
*φ*_*p*_: Fractional reduction of permease activity in the presence of glucose, 1 *- E*_*p*_/*E*_*p,t*_
*k*_1_, *k*_2_: Proportionality constants
*K*_*d*_: Dissociation constant for complex 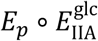
OD_600_: Optical density at 600 nm
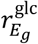: Sp. rate of β-galactosidase synthesis in the presence of glucose + glycerol + TMG
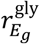: Sp. rate of β-galactosidase synthesis in the presence of glycerol + TMG
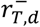: Specific rate of diffusive efflux of TMG
*T*: Intracellular TMG concentration
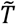: Steady-state intracellular TMG concentration
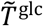: Steady-state intracellular TMG concentration in the presence of glucose
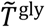: Steady-state intracellular TMG concentration in the presence of glucose
*T*_*e*_: Extracellular TMG concentration
*µ*^glc^: Specific growth rate of *E. coli* cells on glucose
*µ*^gly^: Specific growth rate of *E. coli* cells on glycerol
*X*: Biomass or cell density, gdw l^-1^

