## Supplemental for "Deconstructing glucose-mediated catabolite repression of the *lac* operon of *Escherichia coli*: I. Inducer exclusion, by itself, cannot account for the repression"

### Supplement

#### S1. Steady state intracellular TMG levels in the presence of chloramphenicol

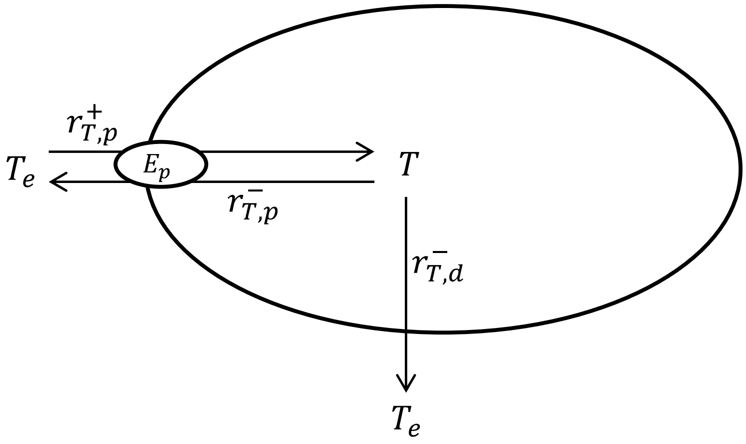


Fig. S1 Kinetic scheme for accumulation and expulsion of intracellular TMG. The notation is explained in the text.

In the presence of chloramphenicol, there is no protein synthesis, and hence, no growth. The lactose enzymes are therefore neither synthesized nor diluted (by growth), and remain frozen at the very same values they had before the addition of chloramphenicol. If such cells are exposed to extracellular TMG, they accumulate TMG until steady state is reached. We show below that the steady state intracellular TMG concentration is proportional to the prevailing specific permease activity provided we (a) correct for TMG influx due to diffusion and (b) eliminate TMG efflux mediated by the permease.

Fig. S1 shows the kinetic scheme of the model. Extracellular TMG, denoted $T_e$, is actively transported into the cell by the permease, denoted $E_p$. Intracellular TMG, denoted $T$, is passively exported to the medium by diffusion and by permease-mediated efflux.

Let $T$ denote the concentration of intracellular TMG; let $r_{T,p}^{+}$ and $r_{T,p}^{-}$ denote the specific rates of permease-mediated influx and efflux, respectively; and let $r_{T,d}^{-}$ denote the specific rate of diffusive efflux. The mass balance for intracellular TMG is given by the equation

$$\frac{dT}{dt} = r_{T,p}^{+} - r_{T,p}^{-} - r_{T,d}^{-}, \qquad \text{S1}$$

where the dilution term has been ignored because there is no growth in the presence of chloramphenicol.

It remains to specify the kinetics. We assume that:

1. The specific rates of permease-mediated influx and efflux follow Michaelis-Menten kinetics, i.e.,

   $$r_{T,p}^{+} = k_{T,p}^{+} E_p \frac{T_e}{K_{T,p}^{+} + T_e}, \qquad \text{S2}$$

   $$r_{T,p}^{-} = k_{T,p}^{-} E_p \frac{T}{K_{T,p}^{-} + T}, \qquad \text{S3}$$

   where $K_{T,p}^{-} \gg K_{T,p}^{+}$.

2. The specific rate of diffusive efflux is given by the equation

   $$r_{T,d}^{-} = k_{T,d}^{-}(T - T_e). \qquad \text{S4}$$

It follows that the steady state intracellular TMG levels are given by the equation

$$0 = \frac{dT}{dt} = k_{T,p}^{+} E_p \frac{T_e}{K_{T,p}^{+} + T_e} - k_{T,p}^{-} E_p \frac{T}{K_{T,p}^{-} + T} - k_{T,d}^{-}(T - T_e). \qquad \text{S5}$$

In inducer exclusion experiments, one measures the steady state intracellular TMG concentration $\tilde{T}$ at fixed specific permease activity $E_p$ and extracellular TMG concentration $T_e$. In what follows, we determine $\tilde{T}$ at various $E_p$ and $T_e$ for three different cases, namely cryptic cells, and wild-type cells exposed to small and large $T_e$.

**Case 1**: Cryptic strains, i.e. $E_p = 0$.

Under these conditions, the steady state intracellular TMG concentration $\tilde{T}_{\text{cryptic}}$ is given by the equation

$$0 = \frac{dT}{dt} = -k_{T,d}^{-}\left(\tilde{T}_{\text{cryptic}} - T_e\right) \quad \Rightarrow \tilde{T}_{\text{cryptic}} = T_e, \qquad \text{S6}$$

i.e., the steady state intracellular TMG level in a cryptic strain equals the concentration of extracellular TMG.

**Case 2**: Small extracellular TMG concentration $T_e$

If the extracellular TMG concentration is sufficiently small, the corresponding steady state intracellular TMG concentration is small compared to $K_{T,p}^{-}$, the saturation constant for permease-mediated efflux. Consequently, permease-mediated efflux is negligible compared to permease-mediated influx, and the steady state intracellular TMG level is given by the equation

$$0 = \frac{dT}{dt} \approx k_{T,p}^{+} E_p \frac{T_e}{K_{T,p}^{+} + T_e} - k_{T,d}^{-}\left(\tilde{T} - T_e\right), \qquad \text{S7}$$

which implies that

$$\tilde{T} - T_e = \tilde{T} - \tilde{T}_{\text{cryptic}} = \left(\frac{k_{T,p}^{+}}{k_{T,d}^{-}} \frac{T_e}{K_{T,p}^{+} + T_e}\right) E_p. \qquad \text{S8}$$

Thus, the steady state intracellular TMG concentration, suitably corrected for the accumulation $\tilde{T}_{\text{cryptic}} = T_e$ due to diffusion, is proportional to the specific permease activity (compare the lowest dashed curve with the other curves in Fig. S2). Moreover, the constant of proportionality increases with $T_e$, i.e., the higher the concentration of extracellular TMG used in the experiment, the faster the corrected concentration $\tilde{T} - \tilde{T}_{\text{cryptic}}$ rises with $E_p$.

Although $\tilde{T} - T_{\text{cryptic}}$ increases linearly with $E_p$, its variation with the induction level $E_g$ can be linear or non-linear depending on the absence or presence of glucose. In the absence of glucose, $E_p$ is proportional to $E_g$ because the permease and β-galactosidase are synthesized coordinately. However, in the presence of glucose, $E_p$ is not proportional to $E_g$ because a fraction of the permease molecules is inactivated by inducer exclusion. Since this fraction decreases with the induction level $E_g$, the ratio $E_p/E_g$ increases with $E_g$.

**Case 3**: Large extracellular TMG concentration $T_e$

It follows from the analysis above that if $T_e$ is sufficiently large, the steady state intracellular TMG concentration in induced cells (with large $E_p$) can become comparable to $K_{T,p}^-$. Under these conditions, permease-mediated efflux cannot be ignored, and the steady state concentration intracellular TMG level satisfies the equation

$$0 = \frac{dT}{dt} = k_{T,p}^+ E_p \frac{T_e}{K_{T,p}^+ + T_e} - k_{T,p}^- E_p \frac{\tilde{T}}{K_{T,p}^- + \tilde{T}} - k_{T,d}^-(\tilde{T} - T_e), \qquad \text{S9}$$

which can be solved for $E_p$ in terms of $\tilde{T}$ to get

$$E_p = \frac{k_{T,d}^-(\tilde{T} - T_e)}{k_{T,p}^+ \frac{T_e}{K_{T,p}^+ + T_e} - k_{T,p}^- \frac{\tilde{T}}{K_{T,p}^- + \tilde{T}}} \qquad \text{S10}$$

It follows that the variation of $\tilde{T}$ with $E_p$ and $T_e$ has the form shown in Fig. S2. Indeed, the above equation implies that $E_p$ increases with $\tilde{T}$. Moreover, $E_p = 0$ when $\tilde{T} = T_e$, and $E_p$ becomes infinite (i.e., $\tilde{T}$ becomes independent of $E_p$) when

$$k_{T,p}^- \frac{\tilde{T}}{K_{T,p}^- + \tilde{T}} = k_{T.p}^+ \frac{T_e}{K_{T,p}^+ + T_e} \qquad \text{S11}$$

The last two results have simple biological interpretations. First, $\tilde{T}$ attains the positive value $T_e$ even when $E_p = 0$ because diffusion ensures that TMG accumulates even in the absence of permease. Second, $\tilde{T}$ becomes independent of $E_p$ at sufficiently large $E_p$ because under these conditions, permease-mediated influx is balanced entirely by permease-mediated efflux, and the effect of permease cancels out. Consequently, if one performs the inducer exclusion experiment with large $T_e$, the steady state intracellular TMG concentrations obtained in the presence of glycerol and glucose may be quite similar even though the corresponding permease activities are quite different.

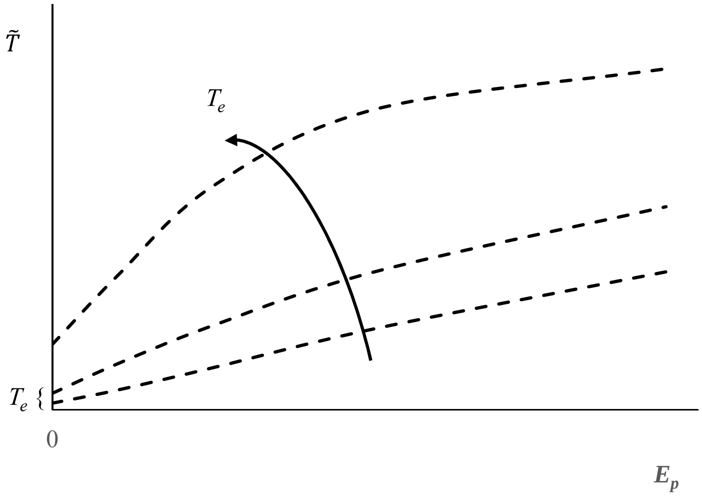


Fig. S2 Predicted variation of the steady state intracellular concentration of TMG $\tilde{T}$ with the specific permease activity $E_p$ at various fixed extracellular TMG concentrations $T_e$. Figure is not to scale.

**S2. Intracellular induction and TMG levels in the absence of chloramphenicol**

In the absence of chloramphenicol, there is protein synthesis and growth. In particular, the lactose enzymes are not only synthesized, but also diluted due to growth. Hence, the evolution of the induction level, as measured by the instantaneous specific β-galactosidase activity $E_g$, is therefore governed by the equation

$$\frac{dE_g}{dt} = r_{E_g}(T, C) - \mu E_g, \qquad \text{S12}$$

where $T$ and $C$ denote the prevailing concentrations of intracellular TMG and cAMP; and $r_{E_g}(T, C)$, $\mu E_g$ denote the specific rates of β-galactosidase synthesis and dilution by growth. The evolution of the intracellular TMG level is now governed by the equation

$$\frac{dT}{dt} = r_{T,p}^{+} - r_{T,p}^{-} - r_{T,d} - \mu T, \qquad \text{S13}$$

which is identical to Eq. S1 except for the appearance of the dilution term $\mu T$. Now, the induction level $E_g$ changes significantly on a time scale of 1 h, whereas the intracellular TMG level $T$ changes significantly on a time scale of 1 min (Fig. S3). It follows that the dilution term is negligible, and $T$ reaches quasi-steady state within 5 min after which its concentration, called the *quasi-steady state* concentration, satisfies the equation

$$0 = \frac{dT}{dt} \approx r_{T,p}^{+} - r_{T,p}^{-} - r_{T,d}. \qquad \text{S14}$$

This equation is identical to Eqs. S1 and S5 that provide $\tilde{T}(E_p, T_e)$, the steady state intracellular TMG level at any given permease activity $E_p$ and extracellular TMG concentration $T_e$. It follows that the quasi-steady state and steady state concentrations of intracellular TMG have the very same functional dependence on the specific permease activity $E_p$ and the extracellular TMG level $T_e$.

We have also shown that the specific permease activity $E_p$ is a function of the induction level $E_g$, which increases linearly in the presence of glycerol, and non-linearly in the presence of glucose. It follows that the steady state concentration of intracellular TMG at any given induction level $E_g$ and extracellular concentration $T_e$ is $\tilde{T}(E_g, T_e) = \tilde{T}(E_p(E_g), T_e)$, and the quasi-steady state concentration of intracellular TMG also has the very same functional dependence on the induction level $E_g$ and the extracellular TMG level $T_e$.

Although the quasi-steady state and steady state concentrations of intracellular TMG have the same functional dependence on $E_g$ and $T_e$, the $\tilde{T}$ vs. $E_g$ data in Figs. Fig. 2 and Fig. 5D are expected to be similar, but not identical, because we used different extracellular TMG concentrations in the two experiments (2 and 100 μM for the data in Figs. Fig. 2 and Fig. 5D, respectively).

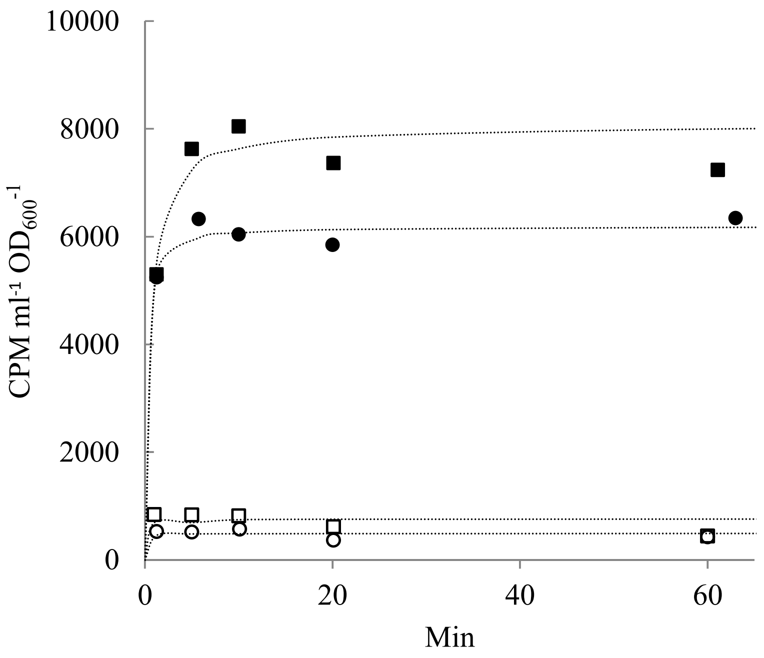


Fig. S3 In inducer exclusion experiments, steady state is achieved within 5 minutes of exposure to [$^{14}$C]TMG. Wild-type cells were induced to varying levels and accumulation of TMG was followed with time. The squares and circles show the data for cells pre-grown on glycerol and glucose. The open and closed symbols refer to uninduced and fully induced cells.
